# Photocurrent production from cherries in a bio-electrochemical cell

**DOI:** 10.1101/2023.01.13.524022

**Authors:** Yaniv Shlosberg

## Abstract

In recent years, big efforts are done to develop new clean energy technologies that will supply the increasing energy demand without contaminating the environment. One of the approaches is the utilization of live organisms as electron donors in bio-electrochemical cells. Photosynthetic organisms may apply for photocurrent generation, releasing NADPH molecules that are formed in the photosynthetic pathway. In this work, we show for the first time that photocurrent can be harvested directly from a cherry fruit associated with a bio-electrochemical cell. Furthermore, we apply electrochemical and spectroscopic methods to show that NADH in the fruit plays a major role in electric current production.

## Introduction

The concern about climate change and the increasing global energy demand has urged scientists to invent creative clean energy technologies to replace the traditional use of fossil fuels. One of the approaches is the utilization of live bacterial cells as electron donors in microbial fuel cells (MFCs)[1]. The electron transfer between the bacteria and the anode may be directly conducted by conductive protein complexes such as metal respiratory complexes[2,3] or *pili* [4–7]. Among the bacterial species that perform high direct exo-electrogenic activity are *Shewanella oneidensis* [5] and *Geobacter. Sulfurreducens* [8]. The electrons can also be transferred by electron mediators that are natively released by the cells such as Quinones and Phenazines. Current enhancement can be done by the addition of artificial electron mediators such as cystine, neutral red, thionin, sulfides, ferric chelated complexes, quinones, phenazines, and humic acids [9–14]. Rather than bacteria, live yeasts may also apply as electron donors in bio-electrochemical cells (BECs)[15]. While yeast consists of reducing molecules, their extracts which is used as a main ingredient in bacterial cultivation media can also apply to produce current and photocurrent in BECs[16]. The current production in BECs may also be catalyzed by the enzymatic activity of [17–20]. An evolution of MFCs is the bio-photo electrochemical cells (BPECs) that use photosynthetic organisms to generate photocurrent. By far, most BPECs studies were based on photosynthetic microorganisms such as cyanobacteria[21–24] and microalgae[25,26]. A unique configuration of a BPEC uses a biofilm in which the cells are arranged at a higher density than a suspension at the surface of the electrode, enhancing the photocurrent production[27]. This approach was also successfully implemented in non-photosynthetic MFCs[5,28,29]. Recent studies showed that BPECs are not limited to photosynthetic microorganisms and can be based on seaweeds[30] and plants [31–33]. A unique class of plants is succulents whose thick cuticle may apply as the envelope of the BPEC, and its inner conductive gel as a native electrolyte[34]. The photocurrent production from seaweeds and plants is about 1000 times greater than reported for micro-organisms[35]. Plants’ roots secrete reducing molecules[36,37] that can also be utilized for electrical current production in BECs[38]. Rather than direct electricity production, the degradation of roots in the soil releases organic compounds that can be used to feed bacteria in soil MFCs[39]. Recently, the variety of organisms’ families that can be used in BECs was broadened to marine animals, showing that *Nematostella vectensis* and *Artemia Salina* can also apply as an electron source [40].

In this work, we broaden the classes of organisms that can be used in BPECs showing for the first time that photocurrent can be harvested directly from a cherry fruit.

## Materials and methods

### Materials

All chemicals were purchased from Merck. Cherries were purchased from Smart and Final store, Goleta, California.

### Fluorescence measurements

Fluorescence measurements were conducted using a Cary Eclipse fluorimeter (Varian) with excitation and emission slits bands of 5 nm, applying a voltage of 700 V on the photomultiplier detector.

### Chronoamperometry measurements

Chronoamperometry measurements were done using palmsens4 potentiostat with screen printed electrodes (Basi), with a graphite working electrode (1mm in diameter), graphite counter electrode, and silver coated with a silver chloride reference electrode. Illumination of the cherry was conducted from the top using a white LED. The light intensity at the electrode height (without the cherry) was determined to be (1500W / m^2^).

### Cyclic voltammetry measurements

Cyclic voltammetry was done using palmsens4 potentiostat with screen-printed electrodes (Basi), with a graphite working electrode (1mm in diameter), graphite counter electrode, and silver coated with a silver chloride reference electrode. Cyclic voltammetry was measured in the range of 0 – 1.2 V, and a scan rate of 0.1 V/s.

## Results and Discussion

### Identification of redox-active molecules in a cherry

Previous studies of BPECs based in succulents[34], reported the advantage of using a bulk-enclosed leaf structure which can apply as an envelope for a BEC, and its inner gel content as a rich electrolyte that also consists of reducing molecules. We wished to study the redox activity of the inner content of a cherry. A cherry was squashed and 50 μL of its inner juice was placed on screen printed electrodes (SPE), with a graphite working electrode (1mm in diameter), graphite counter electrode, and silver coated with a silver chloride reference electrode (Fig. 1a). Cyclic voltammetry (CV) of the juice was measured (0 – 1.2 V, scan rate = 0.1 V/s). A solution of 0.1 M NaCl that should not show any redox peaks in CV measurements applied as a control (Fig. 1b). A positive peak with a maximum of around 0.8 V was obtained for the cherry juice only, which has the voltammetric fingerprint of NADH. This molecule was previously reported to be a major electron mediator in BECs based on various organisms[22,30,40].

**Fig. 1.**
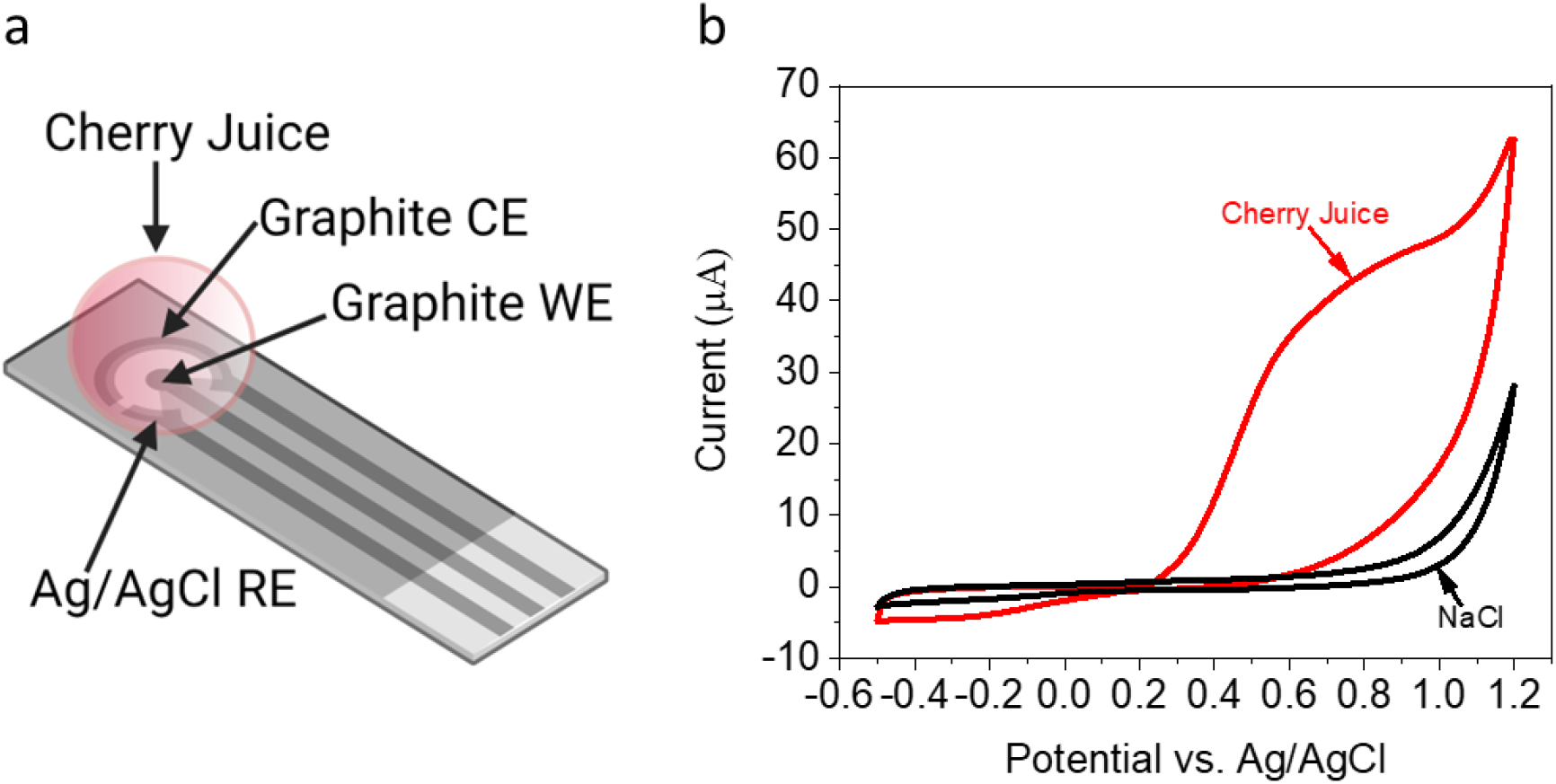
Identification of redox-active molecules in a cherry. A cherry was squashed and 50 μL of its internal solution was placed on SPE. CV of the solution and 0.1M NaCl (as a control) was measured (0 – 1.2 V, scan rate = 0.1 V/s). **a** A schematic description of the system consisted of SPE with graphite working and counter electrodes (WE and CE respectively), and Ag coated with an AgCl Reference electrode (RE). A drop of the cherry’s internal solution (Cherry juice) was placed on top of the electrodes. **b** CV of 0.1M NaCl (black) and Cherry juice (red).

### Identification of NADH in cherry juice by absorption and fluorescence measurements

To support the evidence that the major electron in cherries is NADH, we applied absorption measurements on the cherry’s internal solution. A cherry was squashed, and its juice was diluted by 10 times with water. Absorption spectra of the diluted juice were measured (fig. 3a). The spectra showed peaks around 220, and 280 nm, which corresponds to organic matter[33] and amino acids[41] respectively, and a shoulder at 340 nm which fits the spectral fingerprint of NADH[42]. Although, this shoulder is compatible with the maxima of NADH it may also originate from other unknown molecules that absorb in this range. A more specific method for the detection of NADH is fluorescence[22]. To further strengthen the identification of NADH, fluorescence spectra of the diluted juice were measured (λ_Excitation_ = 340 nm, λ_Emission_ = 400 - 600 nm). The obtained results showed a peak with maxima around 450 nm that correspond to NADH[22] (Fig .2b). This result is in agreement with the absorption (Fig. 2a) and CV measurements (Fig. 1b) supporting the hypothesis that NADH plays a major role in the current generation. Interestingly, this molecule was previously reported to play a key role in mediating electrons in BECs based on different organisms such as cyanobacteria[23], algae[25], and marine animals[40].

**Fig. 2.**
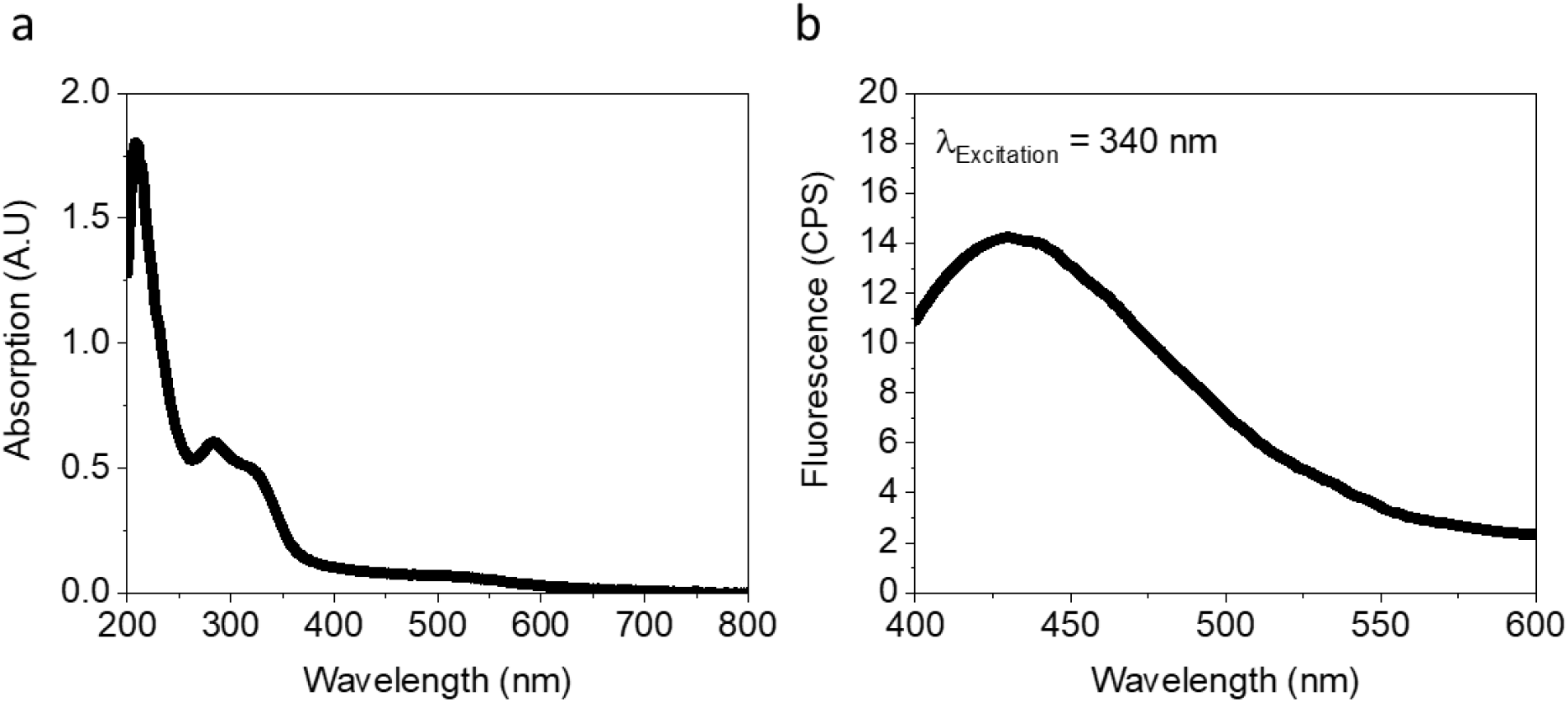
Identification of NADH in cherry juice by absorption and fluorescence measurements. Absorption and fluorescence measurements were conducted for the internal solution of a cherry. The solution was diluted by a factor of 10 to enhance its transparency to enable valid spectroscopic measurements. **a** Absorption spectra of the cherry juice. **b** Fluorescence spectra of the cherry juice (λ_Excitation_ = 340 nm, λ_Emission_ = 400 - 600 nm).

### Direct photocurrent production from a cherry

Based on the observation that cherries consist of a relatively big amount of NADH and other known redox active molecules such as Anthocyanins[43] and Ascorbic acid[44][45]. We wished to explore whether electrical and photoelectrical current can be harvested from a cherry. To do that, SPE was inserted directly into the center of a cherry (horizontally), 5 mm below its top. Illumination was conducted from the top of the cherry by a white LED (Fig. 3a). To measure the light intensity at the electrode surface, the top of the cherry (above the SPE) was cut and placed on top of a light meter. The illumination intensity measured at the electrode surface (below the cherry top) was 1000 W/m^2^. As a control experiment, a drop of 0.1 M NaCl was placed below the top cut of the cherry with a transparent thin nylon sheet between the NaCl solution and the cherry top to prevent mixing of the solutions. Chronoamperometry (CA) measurements were conducted in dark/light intervals of 100 s. The obtained results showed a relatively current density of ∼0.8 μA / cm^2^ for the NaCl that was not influenced by the light illumination. This current may originate from the oxygen evolution reaction that occur at 0.9 V. The CA of the cherry showed a dark current density of ∼1.7 μA / cm^2^ that was enhanced by ∼0.3 μA / cm^2^ in light. Also, the overall trend of the current was increasing over time (Fig. 3b,c). We suggest that the dark current originates mostly from NADH in the cherry while the light dependent current derives from Ascorbic acid whose levels are enhanced in light[45]. The overall increasing trend may indicate an accumulation of Ascorbic acid whose formation rate in light is higher than its oxidation rate at the anode.

**Fig. 2.**
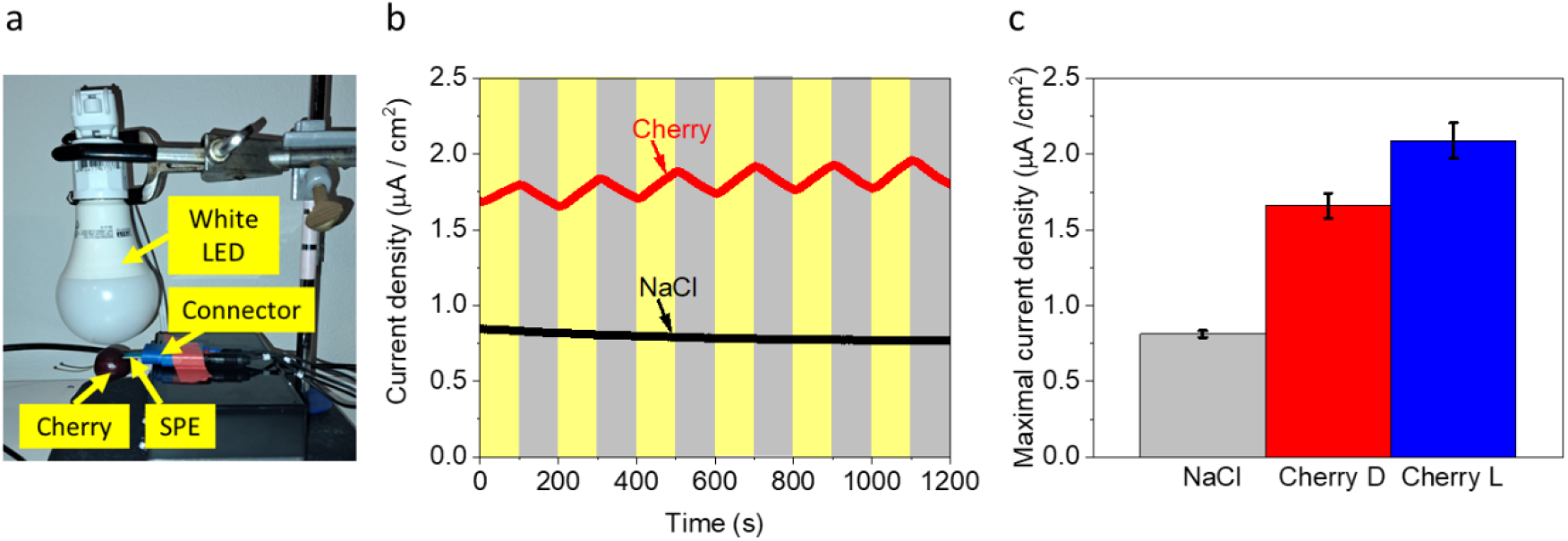
Direct photocurrent production from a Cherry. CA of cherry and 0.1 M NaCl solution (as a control) were measured in dark/light intervals of 100s applying a potential of 0.9 V on the anode. **a** A photo of the system. SPE is inserted into the center of the cherry (5 mm below its top). Illumination is conducted from the top using a white LED. The Intensity at the SPE surface below the cherry was 1000 W/m^2^. The SPE is inserted into a connector that connects it to the potentiostat. Yellow labels with arrows mark the different elements of the system. **b** CA of 0.1 M NaCl (black) and a cherry (red) under dark/light intervals of 100s. Gray and yellow rectangular shapes represent the dark and light during the measurement respectively. **c** Maximal Current density of NaCl (in dark and light, Grey), cherry in dark (Cherry D, red), and cherry in light (cherry L, blue). The error bars represent the standard deviation over 3 independent repetitions.

### Suggesting a mechanism for the electrical current and photocurrent production

Based on the results of this work and previous studies[43–45], we suggest possible mechanisms for the electrical and photoelectrical current generation (Fig. 4). NADH which exists in the internal solution of the cherry, can donate electrons at the anode. Another electron donor is Ascorbic acid whose concentration is elevated upon illumination enhancing photocurrent production. The electron donation at the anode may also originate from anthocyanine which natively applies as an anti-oxidant and can donate electrons. As a pigment, we also suggest a possibility in which upon light illumination, Anthocyanine conducts Förster energy transfer to the anode.

**Fig. 4.**
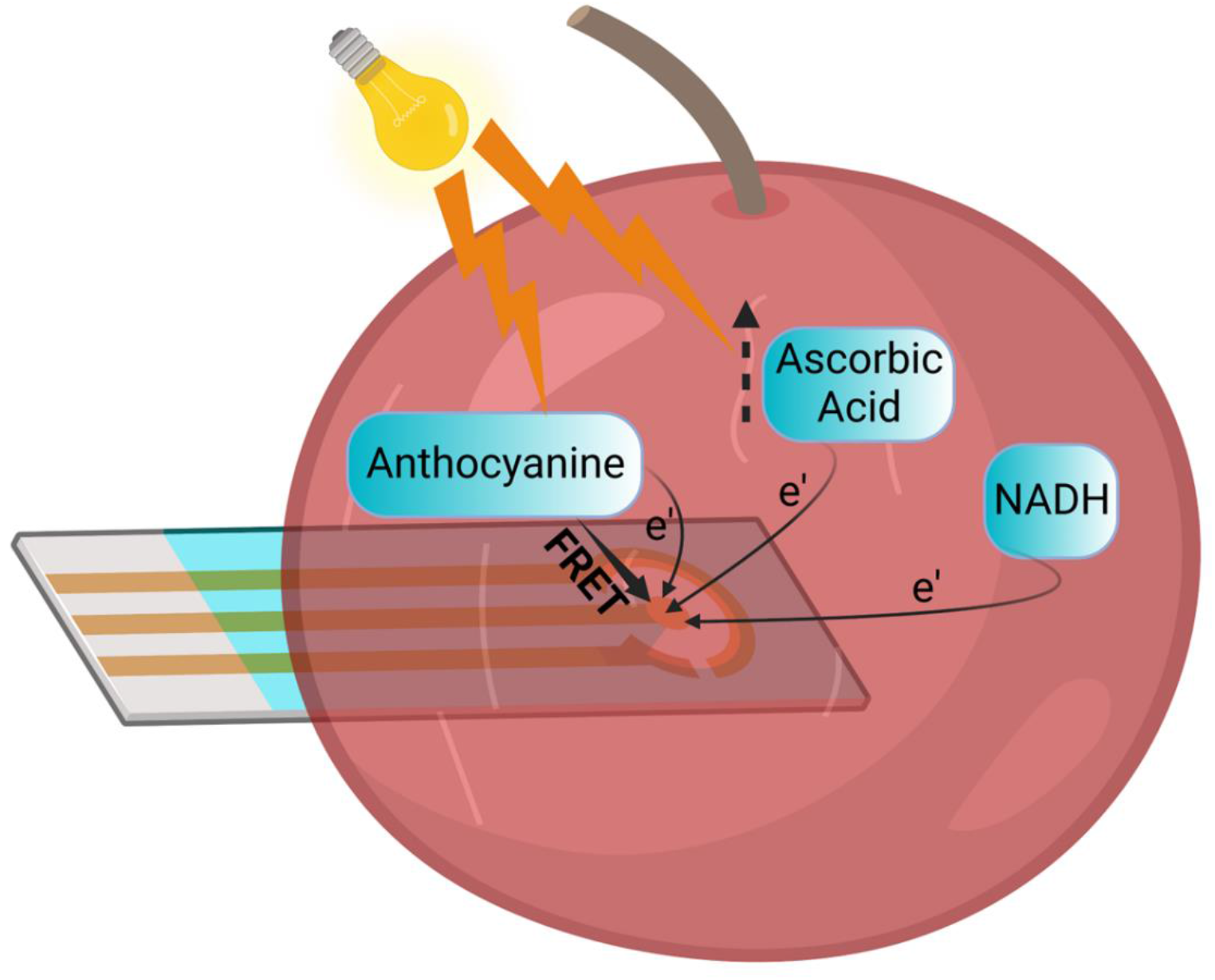
Suggesting a mechanism for the electrical current and photocurrent production. NADH that exists in the cherry can be oxidized at the anode to produce current. The current formation may also originate from Ascorbic acid whose formation is induced by light illumination. Another electron donor may be the pigment Anthocyanine which may also conduct Förster energy transfer in light. a light bulb icon and lightning shapes represent the illumination. Round arrows marked with e’ represent the electron transfer between the electron donors and the anode. A dashed arrow represents the elevation of Ascorbic acid levels that occur in the light. A solid arrow represents the Förster energy transfer from anthocyanine to the anode.

## Conclusions

In this work, we showed for the first time that cherry can be used for a direct photocurrent in a BEC. Also, we show that a major electron donor that plays a role in the current production is NADH. The concept that fruit can be used for photocurrent production may be a base for future clean energy technologies that will exploit fruit species that are not compatible for food or eatable species that are not fresh to generate electricity.

## Acknowledgments

Yaniv Shlosberg is supported by the Otis Williams fellowship. Some of the results reported in this work were obtained using central facilities at the Materials Research Laboratory (MRL). We thank Jaya Nolt for her technical support.

## Author contributions

YS conceived the idea. YS designed the experiments. YS performed the main experiments. YS wrote the paper. YS supervised the entire research project.

